# The role of induced polarization in drug discovery applications

**DOI:** 10.1101/2025.01.14.632933

**Authors:** Ashraf Mohamed, Bernard R. Brooks, Muhamed Amin

## Abstract

Induced polarization plays a pivotal role in ligand-protein binding by enhancing both the specificity and strength of molecular interactions. As a ligand approaches a protein, their respective electronic clouds redistribute in response to each other’s electrostatic fields—a phenomenon governed by induced polarization. The response of a molecule’s electron density to an external field is quantitatively described by its polarizability tensor. In this study, we calculated polarizability tensors for thousands of drug-like molecules from the CHEMBL database, focusing on compounds targeting the Thrombin, Estrogen Receptor alpha, and Phosphodiesterase 5A proteins using Density Functional Theory (DFT). We show that a machine learning model based on atomic hybridization accurately predict the polarizabilities eigenvalues, calculated with DFT. Then, we build a neural network and random forest models to predict the IC50’s based on the same features. The success of these models, despite utilizing a limited number of features, underscores the critical role of induced polarizabilities in determining binding energies.

## Introduction

The concept of induced polarization is pivotal in understanding the mechanisms of biochemical reactions, particularly in drug discovery and molecular interactions.^1-4^ Induced polarization arises when the electron clouds of interacting molecules redistribute in response to external electrostatic fields, enhancing molecular complementarity and stabilizing binding interactions. This phenomenon is central to ligand-protein binding, where the polarization of the electron densities of both ligand and protein contributes to the formation of high-affinity complexes.^5, 6^ By facilitating the precise alignment of electrostatic and van der Waals forces, induced polarization not only strengthens non-covalent interactions but also plays a crucial role in molecular recognition processes, impacting enzyme activity, signal transduction, and drug efficacy. Thus, understanding and quantifying induced polarization provides valuable insights into the molecular basis of key biochemical reactions.

The polarizability tensor, which describes how a molecule’s electron cloud responds to external electric fields, serves as a key metric for quantifying induced polarization. Quantum chemistry methods, such as Density Functional Theory (DFT) and Hartree-Fock calculations, are widely used determine the polarizability tensor with high accuracy.^7-10^ However, these methods are computationally expensive, particularly for large datasets of drug-like molecules. In response to these challenges, machine learning has emerged as a promising alternative for predicting polarizability tensors. By leveraging molecular descriptors, machine learning models can capture relationships between molecular structure and polarizability, offering rapid and scalable predictions.^2, 11-13^ These approaches complement high-accuracy quantum chemistry techniques, enabling the integration of polarizability calculations into large-scale drug discovery workflows.

Molecular descriptors, which encode structural and electronic features of molecules, are essential tools for predicting standard drug discovery properties, such as binding affinities, solubility, and IC50’s values.^14-22^ Common descriptors include molecular weight, topological indices, atom hybridizations, and electrostatic properties. Incorporating polarizability as a descriptor provides additional depth in capturing the molecular response to interactions, particularly for complex environments such as protein binding sites.^23-29^ This inclusion enhances the predictive capability of machine learning models, making them more reliable for assessing molecular performance in dynamic biochemical environments. Furthermore, integrating polarizability into predictive frameworks enables a more comprehensive understanding of ligand-receptor interactions, potentially identifying new therapeutic candidates and accelerating the drug discovery process.

Here, we calculate the polarizability tensors for thousands of molecules from the CHEMBL database using DFT. These molecules are candidates for target proteins Thrombin, Estrogen Receptor alpha, and Phosphodiesterase 5A proteins. Then, we build a machine learning model to predict the eigenvalues of these tensors based on the atomic hybridizations and the eigenvalues of the inertia tensor. Using the same features, we were able to predict the IC50’s for these molecules, which reflects the importance of the induced polarization in computer aided drug discovery.

## Computational Methods

### The DFT calculations

*The DFT calculations* are done with the PSI4 package using B3LYP functional and 6-31G* basis sets.^30^ The structures of the molecules are constructed using the rdkit package based on their SMILES codes. The inertia tensor is calculated based on the atomic masses and coordinates, then the tensor is diagonalized to obtain the eigenvalues.

### Data preparation

The features used in this study are the number of carbons, nitrogens, oxygens, and sulfur in each possible hybridization (sp3, sp2, and sp). The features also include the number of electrons (N) and the number H, Cl, F, Br, and P, since they all have a fixed hybridization in the dataset. Table 1 shows the populations of each hybridization element in each dataset of the different target proteins. To enhance the dataset for each target protein, data augmentation is performed through up-sampling with replacement. To avoid introducing bias training and overfitting, the up-sampling weight is optimized for each model and for each target protein independently.

**Table 1:**
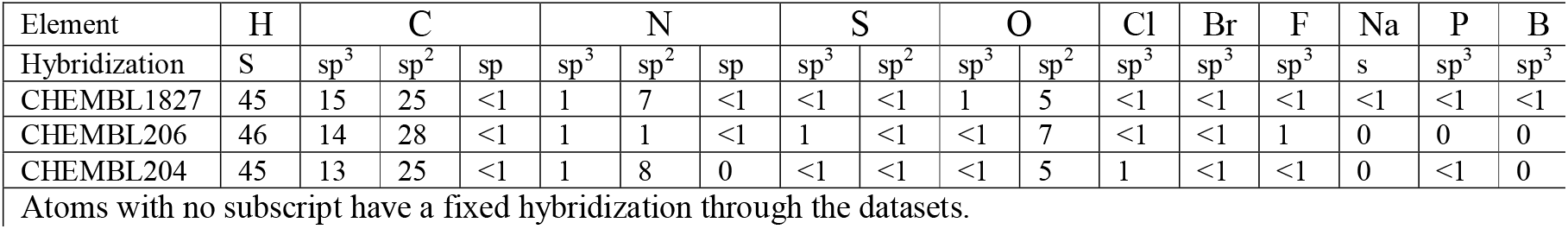
The population percentage (%) of each hybridization component in each dataset.

To ensure the reliability of the dataset, outliers in the IC50’s were removed using the Interquartile Range (IQR) method with a relaxed multiplier. Specifically, the first quartile (Q1) and third quartile (Q3) of the SV distribution were computed, and the IQR was defined as the difference between Q3 and Q1. A relaxed multiplier of 0.6 was applied to extend the acceptable range for SV values, defined as [*Q*1 − 0.6 × *IQR, Q*3 + 0.6 × *IQR*]. Data points outside this range were excluded. This approach balances the need to remove extreme outliers while retaining a broader range of data points for analysis.

### Data splitting

To ensure robust model generalization and minimize potential biases or overfitting, the datasets for the target proteins is split into 80% for training and 20% for validation. Furthermore, to ensure that all the input features contribute equally to the model performance, especially when they are at different scales, the ‘*StandardScaler*’ technique is used. This technique standardizes the features in the dataset by removing the mean and scaling them to unit variance.

### Machine Learning model

The machine learning models are multi-output regressors implemented using the PyTorch library^31^. The models include 24 features in total. Two separate models are trained: one for predicting polarizability and another for predicting the standard value (SV), i.e. the IC50’s. The neural network models used in this study are composed of an input layer, linear layers, batch normalization layers, ReLU activation functions, and an output layer. The input layer accepts a feature vector that represents the input data, such as chemical or structural descriptors. The linear layers perform weighted transformations to extract and refine features, progressively reducing or transforming the dimensionality to better represent the underlying patterns. The batch normalization layers stabilize and accelerate training by normalizing the activations, ensuring consistent distributions across the layers, and helping to prevent internal covariate shifts. The ReLU activation functions introduce nonlinearity into the model, enabling it to capture complex, nonlinear relationships within the data. Finally, the output layer maps the refined features to the desired prediction space, yielding outputs such as the polarizability tensor components (*E*_l_, *E*_2_, *E*_3_) or the standard value (SV) depending on the specific task.

These architectural and training parameters are selected based on an extensive grid search to achieve optimal performance^32^. Figure 1 illustrates the architecture of the two neural network models developed for this study: (a) the model for predicting the polarizability tensor components E1, E2, and E3 and (b) the model for estimating the standard value (SV). Both models follow a similar design with fully connected layers, batch normalization layers, and ReLU activation functions. For the polarizability tensor model (Figure 1a), the input layer accepts a feature vector of size 24. This is followed by a linear layer with 128 neurons, a batch normalization layer, and a ReLU activation function. The next stage involves another linear layer with 128 neurons, another batch normalization layer, and a ReLU activation function. The output layer reduces the dimensionality to three neurons, corresponding to the three components of the polarizability tensor (E1, E2, E3).

**Figure 1.**
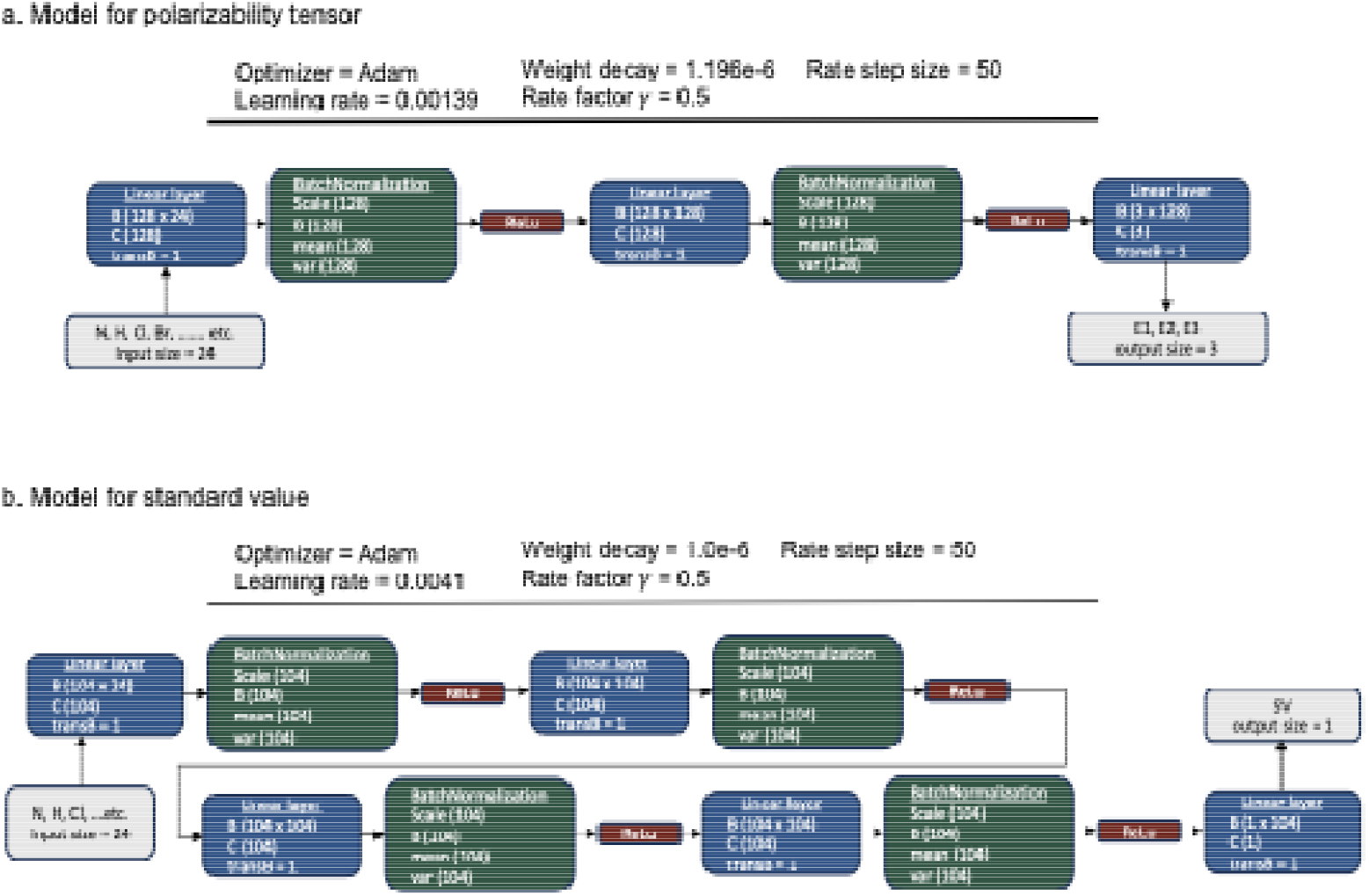
Neural network architectures for (a) polarizability tensor prediction and (b) standard value prediction. Each model consists of fully connected layers, batch normalization layers, and ReLU activation functions. Optimizer settings and hyperparameters, including learning rate, weight decay, and learning rate scheduler parameters, are also detailed. The architectures are optimized for their respective outputs: three components of the polarizability tensor (E1, E2, E3) in (a) and a single standard value (SV) in (b).

In the standard value model (Figure 1b), the architecture is analogous but tailored to a single output. The input layer processes a feature vector of size 24, followed by three similar blocks each of which starts with a linear layer with 104 neurons, a batch normalization layer, and a ReLU activation function. The subsequent linear layer reduces the dimensionality to 104 neurons, followed by batch normalization and ReLU activation. The output layer maps the final representation to a single neuron, which predicts the standard value (SV).

Both models employ the Adam optimizer with specified learning rates and weight decay values. The weight decay acts as a form of regularization, penalizing large weights in the model to reduce overfitting and improve generalization. To ensure effective optimization throughout training, a learning rate scheduler is applied, with a step size of 10 epochs and a reduction factor (*γ*) of 0.5. This scheduler decreases the learning rate by a factor of 0.5 every 10 epochs, enabling the optimizer to take larger steps during the initial phases of training to explore the loss surface and progressively finer steps as training progresses, allowing the model to converge more effectively to a minimum. This approach helps balance faster convergence early on with stability in later stages of training. The training was conducted for 200:500 epochs with a batch size of 128:1024 depending on the target protein and the target variable.

To optimize the model parameters, we employed the *SmoothL1Los* function, also known as Huber Loss. This loss function is particularly suited for regression tasks as it balances the benefits of Mean Squared Error (MSE) and Mean Absolute Error (MAE). For small errors, it behaves like MSE, providing smooth gradients for stable convergence. For larger errors, it transitions to MAE, making the model robust to outliers. Given the presence of noise and potential outliers in our dataset, *SmoothL1Loss* was selected to ensure stable training while mitigating the influence of extreme values.

The training datasets are combined to train the polarizability model. However, for the IC50’s models, each target protein is trained independently due to the significant variation in IC50’s values observed for the same molecule across different target proteins. All models utilize a K-fold cross-validation approach, with the value of K matching the up-sampling weight applied during data augmentation. To ensure unbiased inference, validation samples are neither augmented nor split.

## Results and discussions

Induced polarization plays an important role in protein-ligand interactions. The components of the polarizability tensor quantify the response of the electron density of the molecule to external electric fields. Thus, it is important to include the polarizabilities as a molecular descriptor for accurate protein-ligand binding affinity predictions.^3^ However, the calculations of the molecular polarizabilities require DFT calculations, which are expensive and not feasible to large number of molecules.

An empirical model based on the atomic hybridizations has been previously proposed to provide highly accurate estimates of the average polarizabilities in small molecules^33^ and proteins.^1^ Thus, we build a neural network model to predict the eigenvalues of the polarizability tensor based on the atomic hybridizations. The training and testing datasets are obtained by calculating the polarizability tensors for thousands of molecules, which are drug candidates for the Thrombin, Estrogen Receptor alpha, and Phosphodiesterase 5A target proteins. Furthermore, we added the eigenvalues of the inertia tensor to the features due to the strong correlations between the shape of the molecules and their polarizabilities (Figure 2).^2, 34^

**Figure 2:**
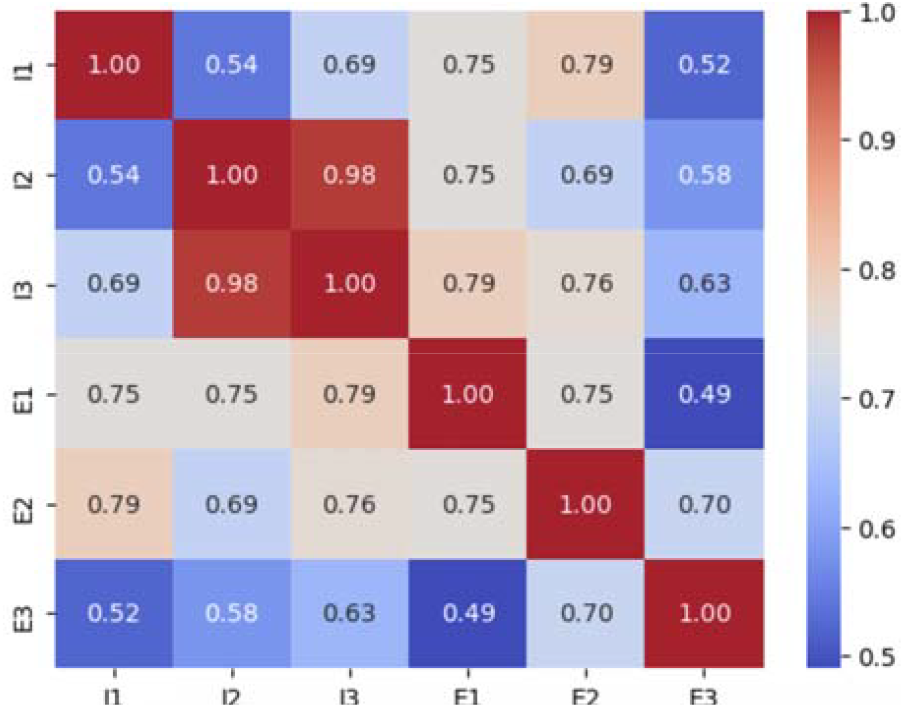
The correlation matrix between the inertia and polarizability eigenvalues. The matrix is constructed using 3210 molecules. The red color indicates a strong correlation, and the blue is for the week correlation.

The model shows a strong correlation between the calculated (using DFT) and the predicted eigenvalues with a correlation coefficient R^2^ of 0.88 ± 0.012 and 0.82 ± and a mean squared error of 0.15 ± 0.011 and 0.18 ± for training and validation datasets respectively (Figure 3). The results are confirmed by cross-validation steps of 5-folds. The contribution of each of the input features is computed by measuring how the prediction changes when a feature is included or excluded. The feature importance analysis, visualized in Figure 4, provides insights into the factors that most strongly influence the model’s predictions. The results show that features related to eigenvalues of the inertia tensor, hydrogen content and the hybridization states of nitrogen and carbon atoms have a substantial impact on the model’s output. These findings suggest that these molecular properties are crucial determinants of the predicted values for the three output variables (E1, E2, and E3). Although the model has only 24 features, it achieves high accuracy in calculating all the polarizability eigenvalues compared to other models, which have a significantly higher number of features.^13^

**Figure 3:**
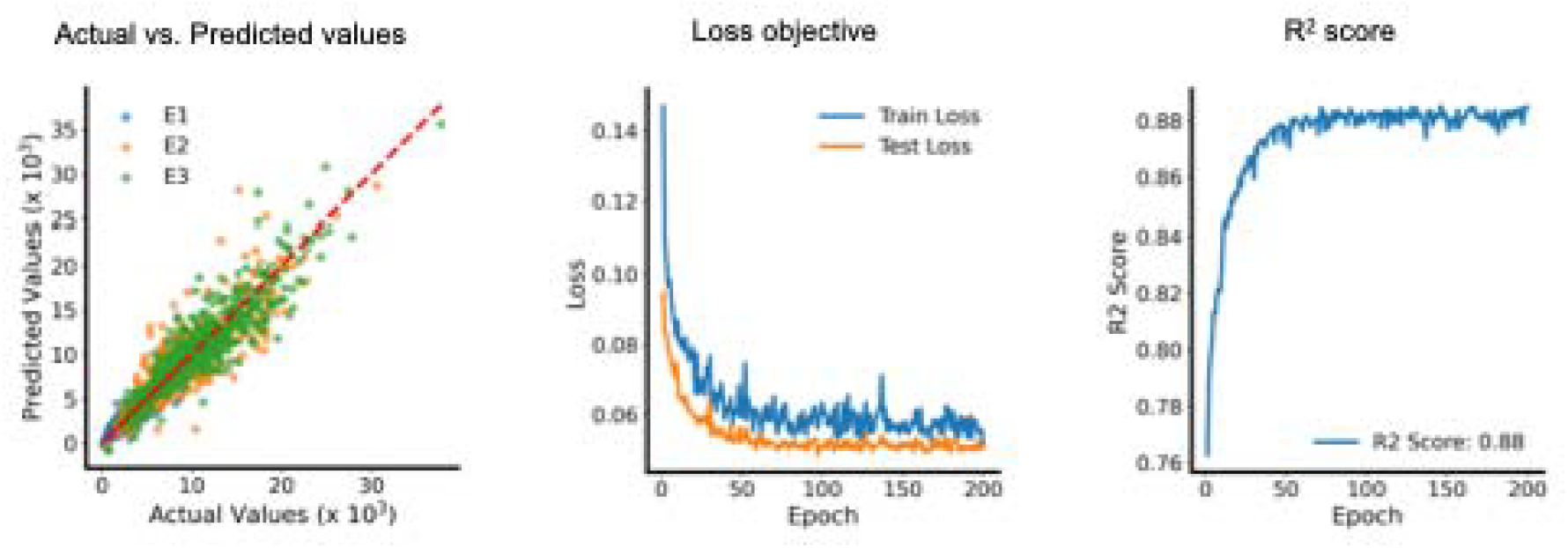
Model performance metrics for polarizability tensor eigenvalues. The dataset includes 3210 points from the three target proteins CHEMBL1827, 204, and 206.

**Figure 4:**
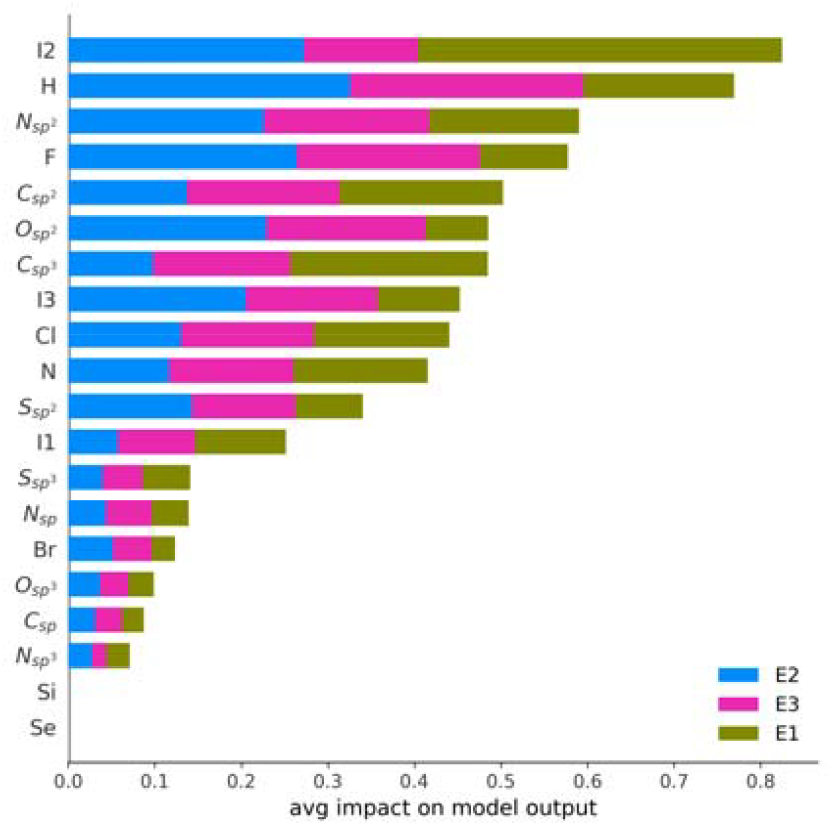
Feature importance analysis. Bar chart showing the average impact of different molecular features on the model’s prediction of three output variables (E1, E2, E3).

The atomic hybridizations characterize the shape of the molecules because they dictate the spatial arrangement of orbitals involved in bonding, which in turn determines the molecule’s geometry. For instance, sp hybridization corresponds to a linear geometry, while sp^2^ and sp^3^ hybridizations result in trigonal planar and tetrahedral geometries, respectively. Consequently, atomic hybridizations can serve as effective shape descriptors, similar to algebraic topology and differential geometry-based descriptors.^35, 36^ Moreover, as shown previously, they are particularly valuable for capturing molecular polarizability, offering a dual-purpose utility in molecular characterization. Thus, we built a neural network model for predicting the IC50’s of three target proteins, Thrombin, Estrogen Receptor alpha, and Phosphodiesterase 5A (CHEMBL206, CHEMBL204, CHEMBL1827) based on the atomic hybridizations of the molecules.

The predicted IC50’s are strongly correlated with the measured values with R^2^ score values from 0.82 to 0.93 when models are trained with K-Fold cross validation for the CHEMBL1827, CHEMBL206, and CHEMBL204 targets respectively (Figure 5) and very low mean square errors (Table 2). Similar to the polarizability model, the feature importance analysis, visualized in Figure 6, provides insights into the factors that most strongly influence the model’s predictions for each SV model separately. The results show that features related to eigenvalues of the inertia tensor, hydrogen content and the hybridization of nitrogen and carbon atoms have a relatively strong impact on the model’s output. These findings suggest that these molecular properties are crucial determinants of the predicted values for the IC50’s.

**Table 2:** The evaluation of the machine learning model for predicting the standard values.

| Target Protein | Neural Networks |  |  |  |  | Random Forest |  |
| --- | --- | --- | --- | --- | --- | --- | --- |
|  | Train R<sup>2</sup> | Train MSE | Val. R<sup>2</sup> | Val. MSE | K | R<sup>2</sup> | MSE |
| ChEMBL1827 | 0.82 ± 0.016 | 0.12 ± 0.014 | 0.79 ± 0.030 | 0.20 ± 0.003 | 3 | 0.66 ± 0.02 | 0.98 ± 0.07 |
| ChEMBL206 | 0.85 ± 0.020 | 0.15 ± 0.030 | 0.74 ± 0.003 | 0.31 ± 0.005 | 10 | 0.66 ± 0.01 | 0.96 ± 0.04 |
| ChEMBL204 | 0.93 ± 0.010 | 0.07 ± 0.001 | 0.86 ± 0.003 | 0.15 ± 0.002 | 5 | 0.54 ± 0.02 | 1.10 ± 0.05 |

The reported metrics are obtained as follows: For the test samples in k-fold cross-validation, Train R<sup>2</sup> and validation R<sup>2</sup> are calculated using the training and validation datasets, respectively. The same approach applies to train MSE and validation MSE. In the Neural Network and Random Forest models, the reported errors are calculated by averaging the performance across the K models from k-folds cross validation. For the validation dataset, the error is determined as the average performance over 100 bootstrapped validation samples.

**Figure 5:**
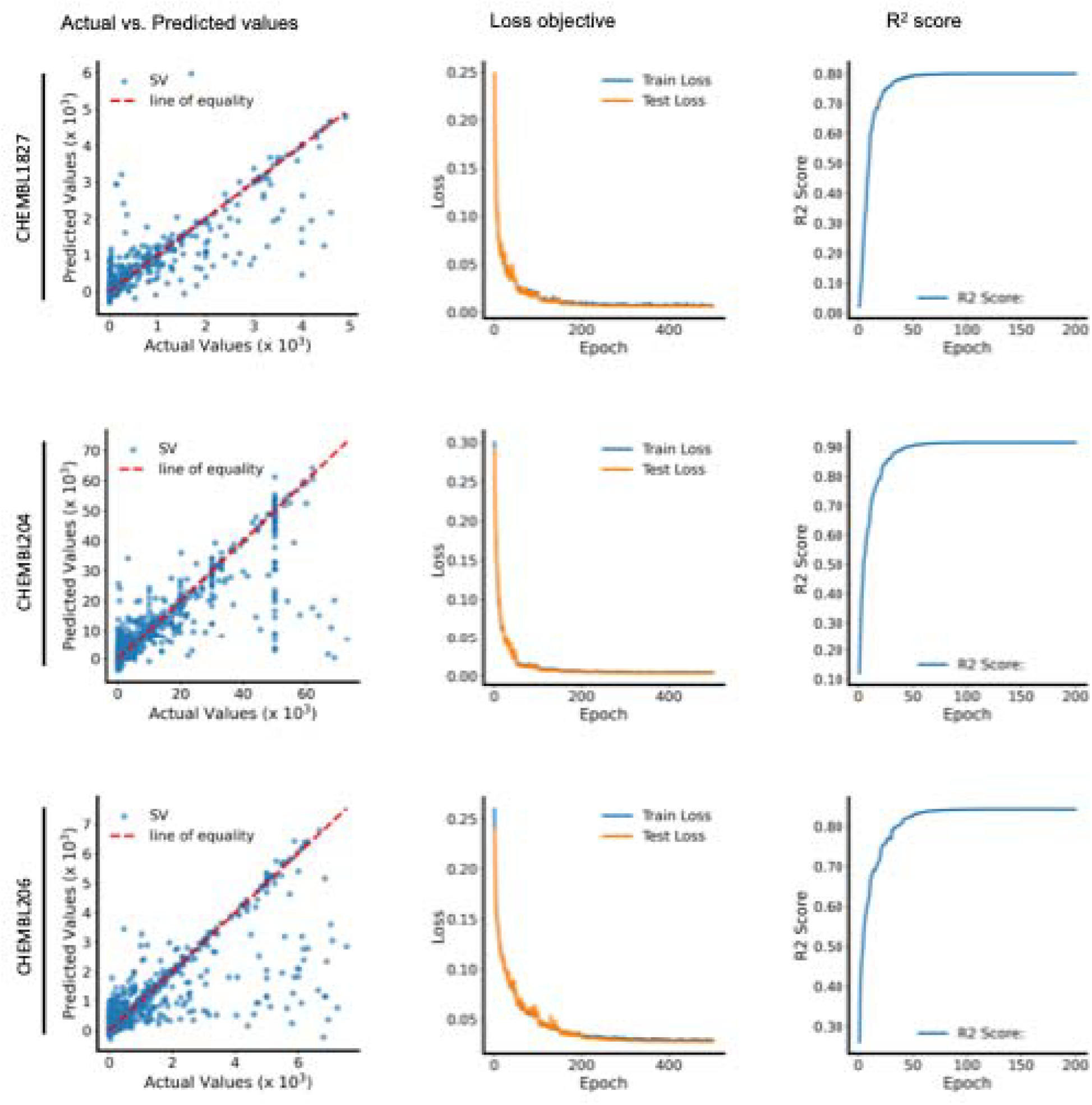
Model performance metrics for IC50’s, comparison between IC50’s predicted vs actual (left), Training and testing loss curves (middle), and R-squared score over epochs when calculated for the test sample (right). Top, middle, bottom are for target proteins CHEMBL1827, 204, and 206 respectively.

**Figure 6:**
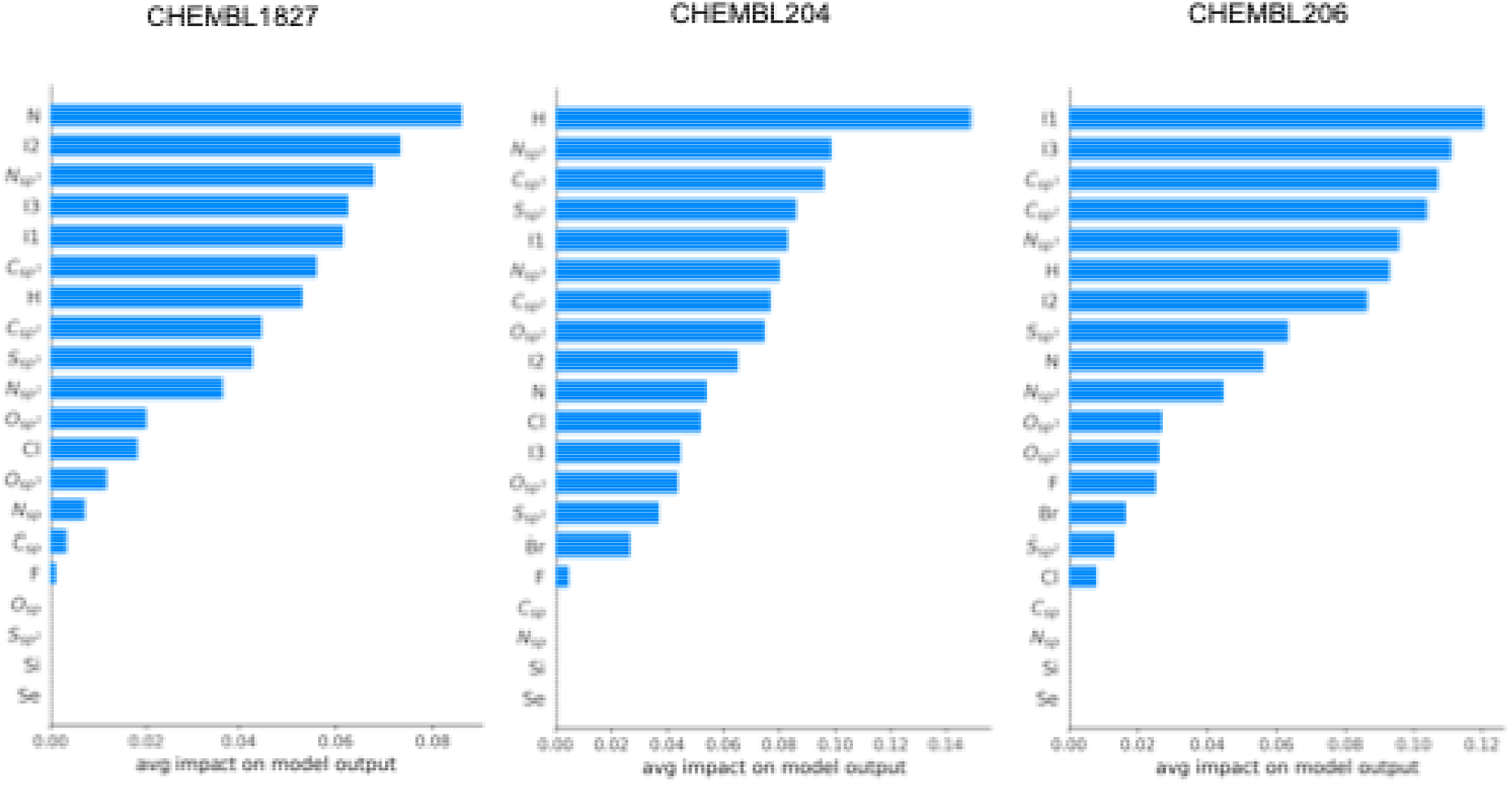
Feature importance analysis. Bar chart showing the average impact of different molecular features on the model’s prediction of the output variable (SV) for the models of the three target proteins CHEMBEL1827, CHEMBEL204 and CHEMBEL206 respectively.

To verify the robustness of our conclusions, we used a simpler model, i.e a random forest regressor (Figure 7). Random forest typically works well “out of the box” with minimal hyperparameter tuning, unlike Neural Networks, which require careful tuning of learning rates, number of layers, neurons, and other parameters. In addition, random forests use an ensemble of decision trees and average their outputs, making them less prone to overfitting. However, because of the large variation in the standard values, the target variable in the random forest model is the log of the IC50’s. Conversely, Neural Networks do not require such transformations. They inherently handle variations through their architecture, particularly by using transformation layers such as batch normalization and activation functions. These layers enable the network to learn patterns directly from the raw IC50’s values without needing explicit log transformations, thus maintaining the original scale of the target variable.

**Figure 7:**
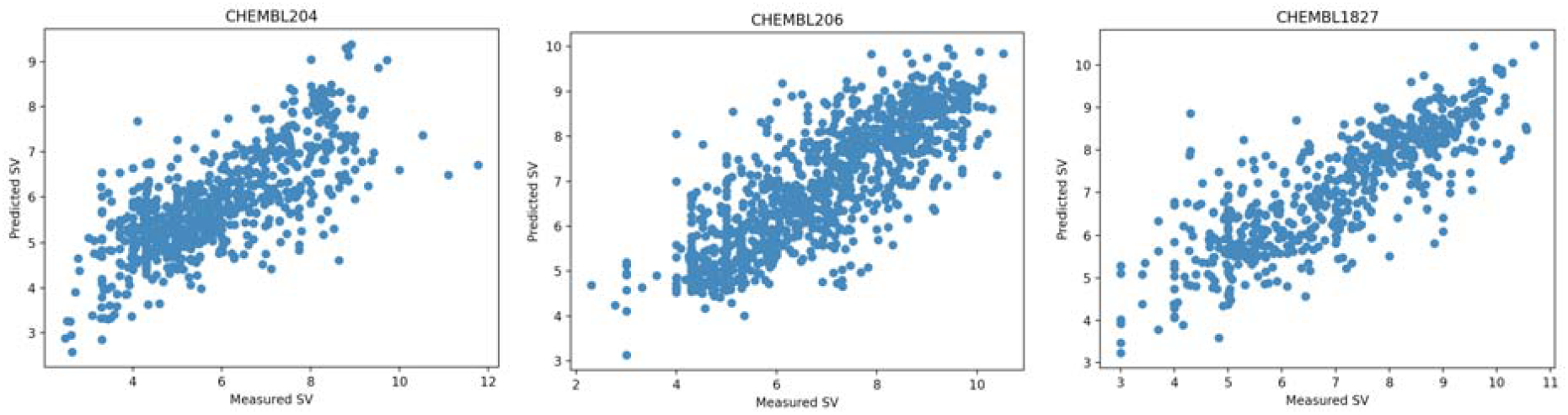
The measured vs. predicted IC50’s for the three target proteins (IDs are on the title) using the random forest regressors. The R2 values are reported in Table 2.

The R^2^’s scores of the predicted vs. experimental IC50’s are 0.66, 0.66 and 0.54 for the CHEMBL1827, CHEMBL206 and CHEMBL204 datasets respectively, which correlates with the R^2^’s obtained from the NN model. The highest MSE value is obtained for the CHEMBL204 dataset. The variation between the predictability of the different target proteins is attributed to the difference in induced polarizability contribution in the binding energies. Additional molecular descriptors may be added to the model to improve the correlation and minimize the error.

## Conclusions

The induced polarization is an important molecular property that should be considered for predicting the target properties of the molecules. Although calculating the polarizability tensors of the molecules using quantum chemistry methods is expensive and inefficient for large datasets, including the atomic hybridizations of the constituent atoms and the inertia principal components is sufficient to account for the molecular polarizabilities. In addition, the atomic hybridizations work as geometric descriptor that include information on the shape of the molecules. Thus, we showed that using a relatively small number of features (27), i.e. the counts of each type of atomic hybridization that exists in the dataset, we were able to obtain accurate predictions for the polarizabilities and the IC50’s for the test sets.

## Supporting information

The polarizabilities calculated with DFT for each molecule

## Data and Software Availability

The CSV files that contain all the data used are provided in the supporting information.

## Notes

### Competing Interest Statement

The authors have declared no competing interest.

